# To design or not to design? Comparison of beetle ultraconserved element probe set utility based on phylogenetic distance, breadth, and method of probe design

**DOI:** 10.1101/2023.01.06.522983

**Authors:** Grey T. Gustafson, Rachel D. Glynn, Andrew E. Z. Short, Sergei Tarasov, Nicole L. Gunter

## Abstract

Tailoring ultraconserved element (UCE) probe set design to focal taxa has been demonstrated to improve locus recovery and phylogenomic inference. However, beyond conducting expensive *in vitro* testing, it remains unclear how best to determine whether an existing UCE probe set is likely to suffice for phylogenomic inference, or if tailored probe design will be desirable. Here we investigate the utility of eight different UCE probe sets for the *in silico* phylogenomic inference of scarabaeoid beetles. Probe sets tested differed in terms of (1) how phylogenetically distant from Scarabaeoidea taxa those used during probe design are, (2) breadth of phylogenetic inference probe set was designed for, and (3) method of probe design. As part of this study, two new UCE probe sets are produced for the beetle family Scarabaeidae and superfamily Hydrophiloidea. We find that, predictably, probe set utility decreases with increasing phylogenetic distance of design taxa from focal taxa, as well as with narrower breadth of phylogenetic inference probes were designed for. We also confirm previous findings regarding ways to optimize UCE probe design. Finally, we make suggestions regarding assessment of need for *de novo* probe design and reinforce previous proposed methods for maximizing UCE probe design to improve phylogenomic inference.

## Introduction

Ultraconserved elements (UCEs) *sensu* Faircloth et al. (2012), have become widely used for phylogenomic analysis of different insect groups including Coleoptera (Baca et al. 2017, Van Dam et al. 2017, Gustafson et al. 2020, Baca et al. 2021, Kobayashi et al. 2021, Bradford et al. 2022, Sota et al. 2022), Diptera (Buenaventura et al. 2021), Hemiptera (Forthman et al. 2019, Kieran et al. 2019, Forthman et al. 2020), Hymenoptera (Blaimer et al. 2015, Faircloth et al. 2015, Branstetter et al. 2017b, Bossert et al. 2019, Cruaud et al. 2019, Zhang et al. 2020, Longino and Branstetter 2021, Pisanty et al. 2022), Isoptera (Hellemans et al. 2022), and Psocodea (Zhang et al. 2019a, Manchola et al. 2022). Their unique features consisting of a highly conserved core with variable flanking regions allow their use in reconstructing both deep- and shallow-level evolutionary relationships (Crawford et al. 2012, Faircloth et al. 2012, Smith et al. 2014, Manthey et al. 2016, Longino and Branstetter 2020). The shorter length of UCE loci relative to other targeted capture approaches (Karin et al. 2020), also facilitates the incorporation of museum specimens into phylogenomic analyses further increasing their utility (Blaimer et al. 2016, McCormack et al. 2016, Ruane and Austin 2017, Van Dam et al. 2017). It is also becoming increasingly evident that UCE loci are primarily exonic in nature in invertebrates, as opposed to vertebrates where intronic loci are predominant (Van Dam et al. 2021). Thus, UCEs have further practicality in their ability to incorporate transcriptomic data, allowing taxa sampled as part of the major 1Kite initiative (https://1kite.org/) to be included in UCE-based phylogenomic studies (Bossert et al. 2019, Baca et al. 2021).

Given their broad phylogenetic viability and increasing use within insect taxa, it is little surprise there has been a proliferation in the number of available UCE probe sets (reviewed in Zhang et al. 2019b, see also Buenaventura et al. 2021, Hellemans et al. 2022, Liu et al. 2022, Van Dam et al. 2022b). Faircloth (2017) produced several publicly available probe sets for Coleoptera, Diptera, Hemiptera, and Lepidoptera to join the original Hymenoptera probe set (Faircloth et al. 2015), all of which were designed to capture loci across entire orders. However, unlike the situation in vertebrates where the original Tetrapod 2.5kv1 and 5kv1 probe sets (Faircloth et al. 2012) provide abundant locus recovery for analysis (i.e., analyzed matrices contained between ∼1,500 – 4,000 UCE loci) across birds (McCormack et al. 2013), mammals (Esselstyn et al. 2017), squamates (Streicher and Wiens 2017), and clades within Anura (Streicher et al. 2018); the insect probe sets do not appear to have the same universal applicability. The original Hymenoptera probe set, hym-v1, developed using two genomes: *Nasonia vitripennis* and *Apis mellifera*, showed decreased locus recovery within the order as phylogenetic distance away from these taxa increased (Faircloth et al. 2015). This led Branstetter et al. (2017a) to refine the existing hym-v1 probes using additional genomic resources, and develop novel probes for newly identified loci across a more diverse swath of hymenopteran genomes, producing the hym-v2 probe set which consist of old and new probes. As part of this new probe design, the inclusion of two ant genomes resulted in probes tailored for use in ants. Implementing the ant-tailored probes resulted in significantly improved locus recovery in taxa for phylogenomic analysis (Branstetter et al. 2017a). Baca et al. (2017) used the Coleoptera 1kv1 probe set for phylogenomic inference of the beetle suborder Adephaga but found locus recovery strongly diminished. Despite its design for use across the entire order Coleoptera, the genomes used in the Coleoptera 1.1kv1 probe design came entirely from the suborder Polyphaga (Fig. 1), given the genomic resources available at the time (Faircloth 2017). Because of this, Gustafson et al. (2019) designed a novel UCE probe set using adephagan genomes, tailored specifically for use in Adephaga (Fig. 1). Like the hym-v2, the Adephaga 2.9kv1 probe set included probes from the original Coleoptera 1.1kv1 design alongside novel tailored probes, and similar to Branstetter et al. (2017a), *in vitro* testing of the tailored probe set revealed increased locus recovery over the original probe set (Gustafson et al. 2020).

**Fig. 1.**
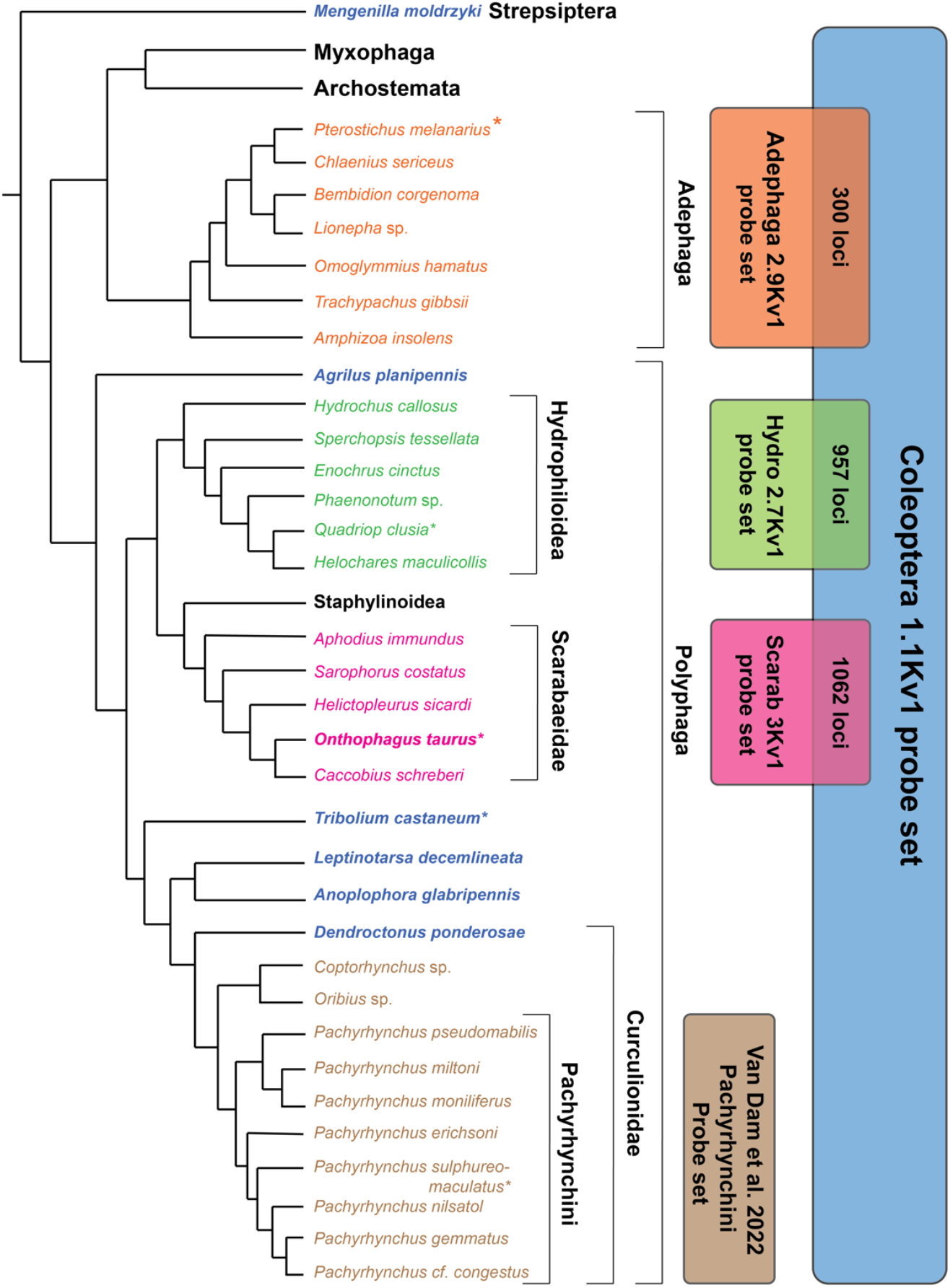
Phylogenetic relationships based on McKenna et al. (2019) among select Coleoptera taxa relevant to, or included in UCE probe design. Color and font of species’ names indicates inclusion in a particular probe design: blue and/or bold font, Coleoptera 1.1kv1 probe design (Faircloth 2017); orange, Adephaga 2.9kv1 (Gustafson et al. 2019); green, Hydro 2.7kv (current study); magenta, Scarab 3kv1 (current study); brown, Pachyrhynchini probe set (Van Dam et al 2022). Asterisk indicates base genome used in final probe design. Boxes associated with the probe set denote the taxonomic breadth probe set was designed to work across. Boxes’ overlap and associated number indicate existence and quantity of loci in common.

While reduced locus recovery in beetles belonging to an entirely different suborder than those used in probe design may be expected, the same is not necessarily true regarding a family of polyphagan beetles that had representation during design. Van Dam et al. (2017) encountered reduced *in vitro* locus recovery in the entimine weevil tribe Eupholini, despite a member of Curculionidae (a scolytine: *Dendroctonus ponderosae*) having been included in the Coleoptera 1.1kv1 probe design (Fig. 1). The estimated divergence time between Entiminae and Scolytinae is ∼90 Ma (Shin et al. 2018), which is comparable to the split between palaeognath birds (ostriches etc.) and the remainder of Aves (Kimball et al. 2019). Given the original tetrapod probe set worked across all of birds and for even deeper divergences within Tetrapoda, this suggests a comparable single universal probe set for all Coleoptera (and likely other insect orders as well) is intractable. The reason for this may have to due with the massive diversity of Coleoptera, including at the genomic level, where varied life histories and diets have produced considerable differences in genome size and complexity through the incorporation and proliferation of horizontally transferred genes (Kirsch et al. 2014, Van Dam et al. 2017, McKenna 2018, McKenna et al. 2019, Lata et al. 2022). Thus, to obtain thousands of loci for analysis, it will likely be necessary to design probe sets tailored to focal clades of beetles. However, a relatively uniform set of homologous loci for use across related taxa within a lineage is desirable for affording cross compatibility between phylogenetic studies, as opposed to the proliferation of isolated probe sets tailored only to the goals of individual studies. Additionally, many research groups may not have the means to generate the genomic resources necessary for producing a novel tailored probe set. Thus, assessing attributes associated with a UCE probe set’s utility for providing robust data for study will aid both in determining need for a novel probe design based on a focal clade and existing probe design, while optimizing odds of *in vitro* success *a priori* when a pre-existing probe set must be selected.

In this study we set forth to investigate the utility of different coleopteran probe sets (Fig. 1) for recovering phylogenetically informative UCE data based on several features including (1) phylogenetic distance from focal taxa, (2) breadth of phylogenetic coverage probe set was designed to provide, and (3) use of methods proposed to optimize probe design. Our goal is to use our findings to provide recommendations for selecting an appropriate existing probe set or identifying if probe design is likely needed. We also set forth to assess previous findings regarding optimization of probe design, to provide recommendations for best practices to produce a tailored probe set. The results of this work have led to the production of two UCE probe sets for the beetle family Scarabaeidae and superfamily Hydrophiloidea.

## Materials and Methods

### Study overview

Our focal taxa are members of Hydrophiloidea and Scarabaeoidea within the beetle suborder Polyphaga. These two lineages have consistently been placed in a clade together along with Staphylinoidea, with recent support for Scarabaeoidea sister to Staphylinoidea (McKenna et al. 2015, Zhang et al. 2018, McKenna et al. 2019). To begin, we generated novel genomic resources for Hydrophiloidea and Scarabaeidae to be used in UCE probe design (Fig. 1). Next during probe design, we experimented with the use of different base genomes, different initial locus identification parameters, and finally merger with pre-existing probes to assess the importance of these factors in probe set design. This resulted in six different novel probe sets: one optimized for base genome, another for initial design parameters, and a final probe set for both Hydrophiloidea and Scarabaeidae that combines fully-optimized, tailored probes and generalized probes taken from the Coleoptera 1.1kv1. Finally, to test the utility of these different probe sets and the publicly available Coleoptera 1.1kv1 and Adephaga 2.9kv1 probe set (Fig. 1), we conducted *in silico* tests using scarabaeoid genomes available through the NCBI database or generated by this study (see Table 1, and supplemental materials Tables S1 & S22). Below we provide detailed methods for each step.

**Table 1.**
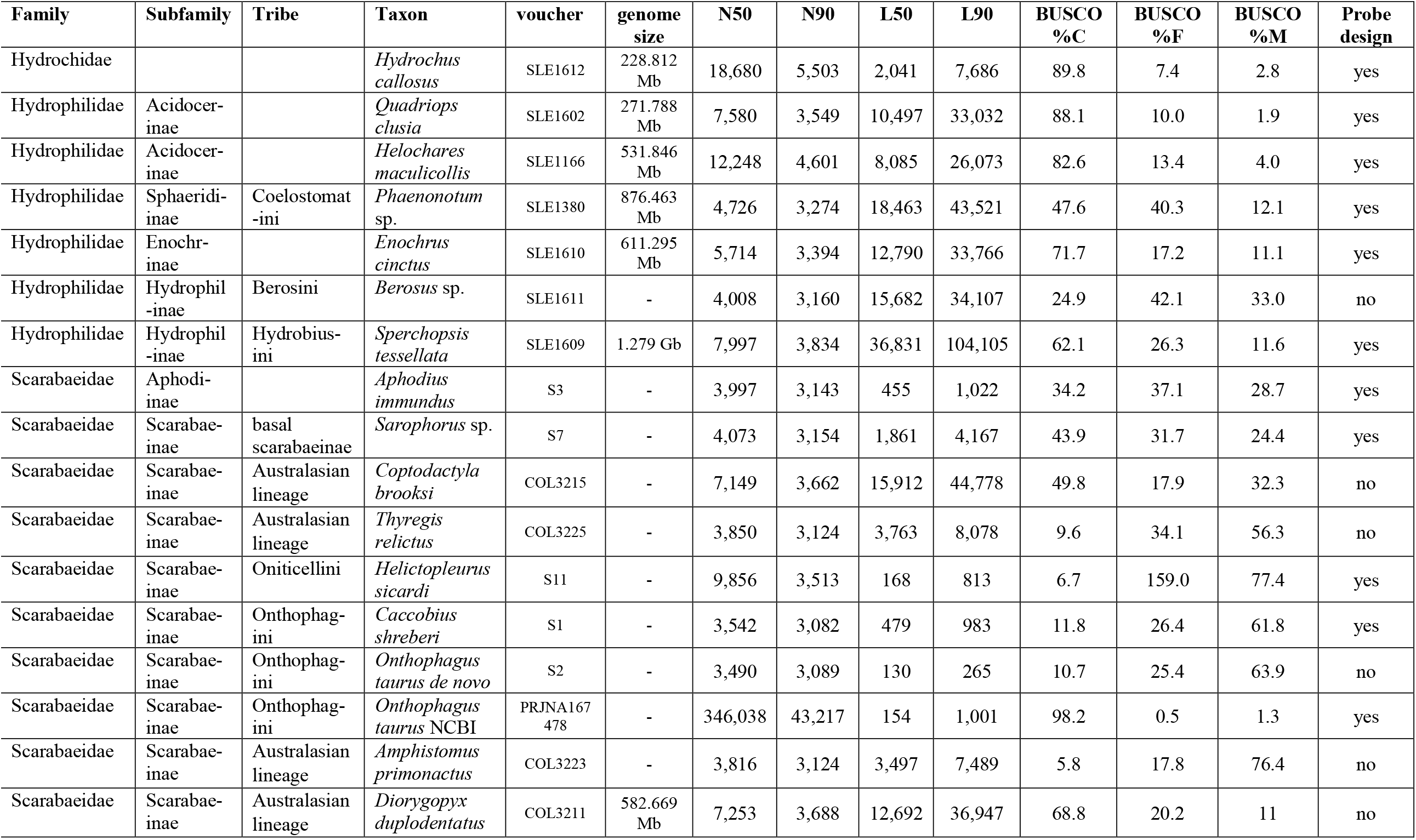
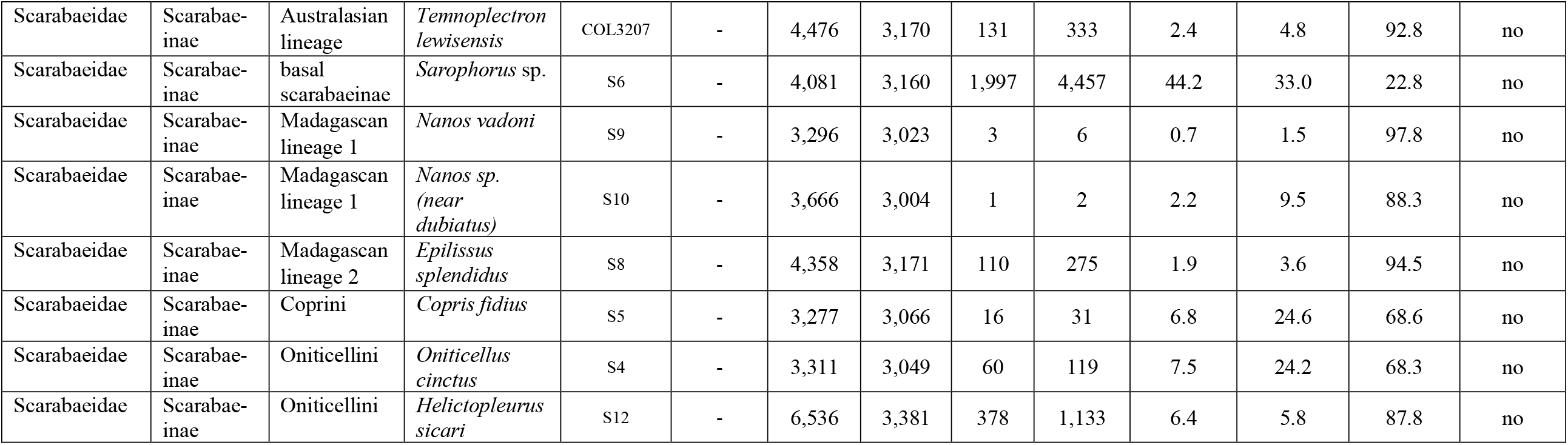
Novel genomic resources generated, their corresponding assembly metrics, and involvement in probe design.

| Family | Subfamily | Tribe | Taxon | voucher | genome size | N50 | N90 | L50 | L90 | BUSCO %C | BUSCO %F | BUSCO %M | Probe design |
| --- | --- | --- | --- | --- | --- | --- | --- | --- | --- | --- | --- | --- | --- |
| Hydrochidae |  |  | *Hydrochus callosus* | SLE1612 | 228.812 Mb | 18,680 | 5,503 | 2,041 | 7,686 | 89.8 | 7.4 | 2.8 | yes |
| Hydrophilidae | Acidocerinae |  | *Quadriops clusia* | SLE1602 | 271.788 Mb | 7,580 | 3,549 | 10,497 | 33,032 | 88.1 | 10.0 | 1.9 | yes |
| Hydrophilidae | Acidocerinae |  | *Helochares maculicollis* | SLE1166 | 531.846 Mb | 12,248 | 4,601 | 8,085 | 26,073 | 82.6 | 13.4 | 4.0 | yes |
| Hydrophilidae | Sphaeridiinae | Coelostomatini | *Phaenonotum* sp. | SLE1380 | 876.463 Mb | 4,726 | 3,274 | 18,463 | 43,521 | 47.6 | 40.3 | 12.1 | yes |
| Hydrophilidae | Enochrinae |  | *Enochrus cinctus* | SLE1610 | 611.295 Mb | 5,714 | 3,394 | 12,790 | 33,766 | 71.7 | 17.2 | 11.1 | yes |
| Hydrophilidae | Hydrophilinae | Berosini | *Berosus* sp. | SLE1611 | - | 4,008 | 3,160 | 15,682 | 34,107 | 24.9 | 42.1 | 33.0 | no |
| Hydrophilidae | Hydrophilinae | Hydrobiusini | *Sperchopsis tessellata* | SLE1609 | 1.279 Gb | 7,997 | 3,834 | 36,831 | 104,105 | 62.1 | 26.3 | 11.6 | yes |
| Scarabaeidae | Aphodiinae |  | *Aphodius immundus* | S3 | - | 3,997 | 3,143 | 455 | 1,022 | 34.2 | 37.1 | 28.7 | yes |
| Scarabaeidae | Scarabaeinae | basal scarabaeinae | *Sarophorus* sp. | S7 | - | 4,073 | 3,154 | 1,861 | 4,167 | 43.9 | 31.7 | 24.4 | yes |
| Scarabaeidae | Scarabaeinae | Australasian lineage | *Coptodactyla brooksi* | COL3215 | - | 7,149 | 3,662 | 15,912 | 44,778 | 49.8 | 17.9 | 32.3 | no |
| Scarabaeidae | Scarabaeinae | Australasian lineage | *Thyregis relictus* | COL3225 | - | 3,850 | 3,124 | 3,763 | 8,078 | 9.6 | 34.1 | 56.3 | no |
| Scarabaeidae | Scarabaeinae | Oniticellini | *Helictopleurus sicardi* | S11 | - | 9,856 | 3,513 | 168 | 813 | 6.7 | 159.0 | 77.4 | yes |
| Scarabaeidae | Scarabaeinae | Onthophagini | *Caccobius shreberi* | S1 | - | 3,542 | 3,082 | 479 | 983 | 11.8 | 26.4 | 61.8 | yes |
| Scarabaeidae | Scarabaeinae | Onthophagini | *Onthophagus taurus de novo* | S2 | - | 3,490 | 3,089 | 130 | 265 | 10.7 | 25.4 | 63.9 | no |
| Scarabaeidae | Scarabaeinae | Onthophagini | *Onthophagus taurus* NCBI | PRJNA167478 | - | 346,038 | 43,217 | 154 | 1,001 | 98.2 | 0.5 | 1.3 | yes |
| Scarabaeidae | Scarabaeinae | Australasian lineage | *Amphistomus primonactus* | COL3223 | - | 3,816 | 3,124 | 3,497 | 7,489 | 5.8 | 17.8 | 76.4 | no |
| Scarabaeidae | Scarabaeinae | Australasian lineage | *Diorygopyx duplodentatus* | COL3211 | 582.669 Mb | 7,253 | 3,688 | 12,692 | 36,947 | 68.8 | 20.2 | 11 | no |
| Scarabaeidae | Scarabaeinae | Australasian lineage | *Temnoplectron lewisensis* | COL3207 | - | 4,476 | 3,170 | 131 | 333 | 2.4 | 4.8 | 92.8 | no |
| Scarabaeidae | Scarabaeinae | basal scarabaeinae | *Sarophorus* sp. | S6 | - | 4,081 | 3,160 | 1,997 | 4,457 | 44.2 | 33.0 | 22.8 | no |
| Scarabaeidae | Scarabaeinae | Madagascan lineage 1 | *Nanos vadoni* | S9 | - | 3,296 | 3,023 | 3 | 6 | 0.7 | 1.5 | 97.8 | no |
| Scarabaeidae | Scarabaeinae | Madagascan lineage 1 | *Nanos* sp. (near *dubiatus*) | S10 | - | 3,666 | 3,004 | 1 | 2 | 2.2 | 9.5 | 88.3 | no |
| Scarabaeidae | Scarabaeinae | Madagascan lineage 2 | *Epilissus splendidus* | S8 | - | 4,358 | 3,171 | 110 | 275 | 1.9 | 3.6 | 94.5 | no |
| Scarabaeidae | Scarabaeinae | Coprini | *Copris fidius* | S5 | - | 3,277 | 3,066 | 16 | 31 | 6.8 | 24.6 | 68.6 | no |
| Scarabaeidae | Scarabaeinae | Oniticellini | *Oniticellus cinctus* | S4 | - | 3,311 | 3,049 | 60 | 119 | 7.5 | 24.2 | 68.3 | no |
| Scarabaeidae | Scarabaeinae | Oniticellini | *Helictopleurus sicari* | S12 | - | 6,536 | 3,381 | 378 | 1,133 | 6.4 | 5.8 | 87.8 | no |

### Novel genomic resource generation

A diverse sampling of taxa from across the Hydrophilidae, representing five of six currently recognized subfamilies and one member from the closely related hydrophiloid family Hydrochidae (Short and Fikáček 2013), were chosen for genomic sequencing and probe design (Table 1). For scarabs, sampling targeted the subfamily Scarabaeinae (13 genera from seven lineages that are equivalent to tribes) and one member of the closely related subfamily Aphodiinae (Table 1). All specimens were preserved in ≥ 95% ethanol and kept frozen at −20° C prior to extraction. Total genomic DNA was extracted using a DNeasy Blood and Tissue Kit (Qiagen, Hilden, Germany) following the manufacturer’s protocols. Extractions were quantified via a Qubit® (Life Technologies, Carlsbad, CA, U.S.A.) system to estimate input DNA for library preparation.

For hydrophiloid taxa, library preparation was done by technicians at RAPiD Genomics LLC (Gainesville, FL, U.S.A.) and the Genomic Sequencing Core at the University of Kansas (KU). There, DNA was mechanically sheared to an average size of 400 base pairs (bp), with libraries constructed by repairing the ends of the sheared fragments followed by the ligation of an adenine residue to the 3’-end of the blunt-end fragments. Then barcoded adapters suited for the Illumina Sequencing platform were ligated to the libraries, with ligated fragments being PCR-amplified using standard cycling protocols (e.g., (Mamanova *et al*., 2010). Genomic sequencing for *Quadriops clusia, Berosus* sp., and *Hydrochus callosus* was done on an Illumina HiSeq 3000 with paired-end 150 bp reads. Genomic sequencing for *Helochares maculicollis, Phaenonotum* sp., *Sperchopsis tessallata*, and *Enochrus cinctus* was done on an Illumina NextSeq 550 with paired-end 150 bp reads. Demultiplexing was also done by techs at RAPiD Genomics LLC and the KU Genomic Sequencing Core. For the scarab taxa used in probe design (non-Australasian lineages, Table 1), a Nextera Flex library (Illumina) was sequenced using an Illumina NextSeq 500 sequencer. For Australasian dung beetles, genomic DNA was sent to SNPsaurus (Eugene, OR) for whole genome genotyping. Following their standard workflow, a Nextera tagmentation kit was used for library generation, samples were subsequently sequenced on a Illumina HiSeq 4000 sequencer with paired-end 150 bp reads, and adapters trimmed with BBDuk (BBMap version 38.41).

Raw demultiplexed Illumina reads were first cleaned using the TRIMMOMATIC (Bolger et al. 2014) with final adapter contamination trimmed using CUTADAPT 1.18 (Martin 2011)) or FASTP (Chen et al. 2018). Raw, cleaned and trimmed reads were quality inspected using FASTQC (Andrews, 2010). For hydrophiloid taxa the best *k*-mer length for *de novo* genome assembly was inferred using KMERGENIE (Chikhi and Medvedev 2014). This program was also used to estimate genome size using *k*-mers from assemblies that were found to be ≥ 50% complete (Table 1). Unless otherwise indicated, genome assembly was done using SPAdes 3.7.1 (Bankevich et al. 2012) under the default *k*-mer settings for 150 bp paired-end reads including the -m 1000 memory option. For *Quadriop clusia, Hydrochus callosus*, and *Helochares maculicollis*, which were identified as having the best *k*-mer assembly size above 77, the -k option was used with the *k*-mer values of 21, 33, 55, 77, 99, 127. For scarab taxa used in probe design (non-Australasian lineages, Table 1) SPARSEASSEMBLER (Ye et al. 2012) was used to assemble low-coverage genomes using forward and reverse reads with the command ‘SparseAssembler g 10 k 31 LD 0 GS 1000000000 NodeCovTh 1 EdgeCovTh 0 1’ as suggested earlier (Kypke 2018, Brunke et al. 2021).

Genomic assemblies were first evaluated using QUAST 5.0.2 and 5.2.0 (Mikheenko et al. 2018) in order to get the following metrics: total contig number, total assembly length, coverage, GC content, N50, N90, L50, L90 (Tables 1, S1). Next BUSCO 3.0.2 (Simão et al. 2015) was used to measure assembly completeness. BUSCO was run using *Tribolium castaneum* as the AUGUSTUS (Stanke et al. 2004) input species and the 2,442 Endopterygota single copy orthologous protein coding reference genes from OrthoDB v9 (Zdobnov et al. 2017). This provided each genome assembly with BUSCO completeness scores (Table 1). Full genome assembly stats are provided in Table S1 in the supplemental materials.

### Probe design: Hydrophiloidea

The hydrophiloid probe design follows the methods described in Gustafson et al. (2019) for the identification of a base genome and optimization of the probe set. Optimal base genome identification can be done either by conducting base genome trials or selecting the taxon with the shortest overall genetic distance from all other taxa (see Gustafson et al. (2019) for more details). A major goal of this design, other than to produce a hydrophiloid probe set, was to test the findings of Gustafson et al. (2019) regarding probe set design optimization.

First, we gathered sequence data for six molecular markers commonly used in phylogenetic studies of the Hydrophilidae (Table S2) (Short and Fikáček 2013) for each genus used in probe design or a sister taxon based on the phylogeny from Short and Fikáček (2013). These loci were generated via Sanger sequencing and are independent of the genomic sequencing conducted here. Additionally, we randomly selected 10 of the genes successfully extracted directly from each genomic assembly that were used to estimate BUSCO completeness scores to generate additional genetic distance measures for comparative purposes.

For these two sets of markers genetic distances among taxa were estimated using four different models: Jukes Cantor (Jukes and Cantor 1969), Tajima Nei (Tajima and Nei 1984), Tamura Nei (Tamura and Nei 1993), and the maximum composite likelihood model (Tamura et al. 2004), as implemented in MEGA ver. 6.06 (Tamura et al. 2013). From the resulting genetic distance measures taxa were ranked from smallest to largest for each model. These rank scores were then averaged across models for each marker. From the average scores for each marker the overall genetic distances ranks were established for each taxon within the two sets of markers (i.e., the six Sanger loci and the 10 BUSCO genes).

Both the Sanger and BUSCO genetic markers identified *Quadriops* as having the shortest average overall genetic distance from all other taxa used in probe design. Therefore, *Quadriops* was selected as the initial base genome for UCE probe design. Design was conducted in PHYLUCE ver. 1.7.1 following the methods outlined in detail in Faircloth (2017) and the generalized workflow discussed in Gustafson et al. (2019). Our temporary baits were initially designed to target putative loci shared among the base taxon and all five other taxa. Following alignment of the temporary baits back to the genomic assemblies, the final probe set was designed to target only loci present in all six taxa. Specific probe design commands and settings are available in the supplemental materials.

After the initial probe design described above was completed, monolithic fasta files were produced using the same methods with each hydrophilid taxon serving as the base genome during probe design. This was done (1) as a base genome test to confirm previous findings that base genome choice significantly alters locus recovery and further test the ability of genetic distance measures alone for selecting an optimal base genome; (2) apply a more stringent paralogy filter to our final probe set using a pipeline consisting of BLAST (Altschul et al. 1990) and R scripts (R Core Team 2022) for comparing monolithic fasta files (i.e., comparing_monolithic_UCE_fasta v0.2 and whittle_uce_probes.R (Alexander 2018 and Gustafson et al. 2019)); and (3) receive further information about locus length and recovery in other taxa given a particular base genome using these R scripts (i.e., select_base_genome.R (Alexander 2018)). Because the *Berosus* genome was largely incomplete (Table 1), it was not included in probe design as another hydrophiline, *Sperchopsis*, was available. This first probe set design represents optimization for base genome.

To test the effect of initial design parameters on final probe design, one additional *de novo* probe design was conducted using the optimal base genome. For this design, putative locus identification for development of temporary baits was done using loci shared between the base genome and just one other taxon (i.e., +1), rather than all other taxa. Probe design then continued identically to that described above, with the exception of swapping base genomes to produce multiple probe sets. Note the paralogy filter was still applied using the same monolithic fastas from the prior probe design but with the new optimal base genome +1 monolithic fasta file replacing the corresponding +6 fasta. This second probe set design represents optimization for initial design parameters.

In addition to the novel UCE loci identified, we wanted to include probes from the Coleoptera 1.1kv1 probe set designed by Faircloth (2017) to produce the final probe set. This will afford cross-compatibility and it has been demonstrated that combining tailored probes alongside the generalized probes of the Coleoptera 1.kv1 can actually improve phylogenetic analysis (Gustafson et al. 2020). When combining probe sets designed in PHYLUCE there are a couple of considerations: (1) there could be homologous loci identified in common between them; (2) the probe naming scheme can lead to homonymous probes that target non-homologous loci (i.e., both probes sets may target a UCE locus 300, but ‘locus 300’ may not be homologous between the two. To combine the probe sets, given these considerations, we generated novel R scripts (available from https://github.com/sergeitarasov/Rename-and-Merge-UCEs) that first BLASTs between monolithic fast a files produced by the two probe sets on focal design taxa to identify homologous loci. Next, probe naming conventions are compared such that new probes targeting homologous loci in common with the original probe set (in this case, Coleoptera 1.1kv1) are renamed to match the original probe set (so probes in both sets target the same locus 300). Finally, the two sets of probes are merged such that duplicate probes from the original probe set are removed to prevent overlap; and homonymous probe names between the two, now target correspondingly homologous UCE loci.

### Probe design: Scarabaeidae

For the Scarabaeidae probe design, we followed all of the methodology above with the exception of performing genetic distance measures, as we encountered difficulties finding 10 BUSCO genes in common across all scarab genomes used in probe design, likely due to their relatively low completeness (Table 1). Additionally, we ultimately wanted the scarab probe design to utilize the available NCBI genome assembly of *Onthophagus taurus*. There were several reasons for this: (1) the assembly is highly complete (Table 1); (2) its annotation will allow all loci targeted by the resultant probe design to be identifiable, thus preventing inclusion of ‘off-target’ probes (see Van Dam et al. 2022a); and (3) it performed relatively well as the base genome.

### UCE locus identification

We used the scripts provided by Van Dam et al. (2021) to identify loci targeted by our probe sets. These scripts require an annotated genome to function. No annotated hydrophiloid genome is currently available, thus we only utilized the *Onthophagus taurus* annotation. To identify hydrophiloid UCE loci using this annotation, we produced *Onthophagus*-based probes by aligning the final Hydrophiloid probe set against the *Onthophagus taurus* genome as ‘temporary baits’, then based on which probes hit, we produced finalized probes using the same methods described above.

### In silico testing

For assessing optimality of *de novo* probe design, i.e., base genome experimentation and relaxed initial design parameters, via PHYLUCE we compared the number of loci extracted by the finalized probe set when applied only to genomes used in the corresponding probe design.

To test the broader phylogenetic utility of the optimized-, final-, and existing UCE probe sets, we downloaded scarabaeoid genomes available through the NCBI website (see Table S22). Two Staphylinoid taxa were included as an outgroup. Scarabaeoid taxa were selected because (1) an abundance of genomic resources relative; and (2) they afford a range of phylogenetic distances from the Coleoptera UCE probe sets (Fig. 1). Specifically, Scarabaeidae is nested within Scarabaeoidea, which in turn together with Staphylinoidea is sister to Hydrophiloidea, and the aforementioned clade is nested inside the suborder Polyphaga, which is sister to the suborder Adephaga. Through the PHYLUCE 1.7.1 pipeline, first probes were aligned to the genomic data and the surrounding sequence information extracted using the command phyluce_probe_slice_sequence_from_genomes with flank value set to 500. Next UCE loci were pulled from these contigs using each UCE probe set with the command phyluce_assembly_match_contigs_to_probes and the minimum identity set at 80 and the minimum coverage 50. Extracted UCE loci were aligned using MAFFT (Katoh and Standley 2013) under the default PHYLUCE wrapper settings, but with the additional command ‘no-trim’ to provide internal trimming. Internal trimming is recommended for analysis of divergences over 50 millions years ago (MYA) (Faircloth 2016), and most splits among our taxa meet this requirement (i.e., *Onthophagus* + *Aphodius* split was estimated ca. 100 MYA in McKenna et al. (2019)). Next, alignments were trimmed with the GBlocks (Castresana 2000, Talavera and Castresana 2007) PHYLUCE wrapper and the commands b1 0.5, b2 0.85, b3 8, b4 10. Finally, an unpartitioned 75% complete concatenated data matrix was produced and the program AMAS (Borowiec 2016) was used to generate summary statistics associated with each data set.

Maximum likelihood analyses were conducted using IQ-TREE ver. 1.6.12 (Nguyen et al. 2015) and the MFP option (Kalyaanamoorthy et al. 2017) to select the optimal evolutionary model using corrected Akaike information criterion scores, and apply it for phylogenetic inference. Nodal support was assessed with 1000 ultrafast bootstraps (Hoang et al. 2018). As relationships among Scarabaeoidea remain unclear, we generated Robinson–Foulds (R-F) distances (Robinson and Foulds 1981) in RAXML ver. 8.2.12 (Stamatakis 2014) to quantitatively assess differences among the resultant phylogenetic trees.

## Results

### Base genome identification and base genome optimized probe sets

The overall average genetic distance measures using both common Sanger gene fragments for phylogenetics (Fig. 2A, Tables S2–S8) and ten randomly selected BUSCO genes (Fig. 2B, Tables S2, S9–S18) both identified *Quadriops clusia* as having the shortest distance from all other taxa used in probe design, implying its identification as the optimal base genome. The Hydrophilioid base genome tests unambiguously confirmed *Quadriops clusia* as being the optimal base genome for probe design in that it resulted in: the highest number of total UCE loci recovered (Fig. 2A,B), the highest number of loci recovered in individual taxa (Table S19), the highest number of paralogy-free loci across all taxa when comparing monolithic fasta files, and the longest locus lengths. Relative to the second best performing base genome, *Helochares*, use of *Quadriops* as the base genome resulted in a probe set that recovers several thousand more loci, representing a 66% increase in total locus recovery (Table S1). We found base genome performance, as assessed by total number of UCE loci recovered during *in silico* testing, to be negatively correlated with average genetic distance rank (Figs. 2A,B). Genome assembly metrics (Table 1) like N90 value (Fig. 2C) and BUSCO %C (Fig. 2D), showed little-to-no correlation with base genome performance. Given the above findings, *Quadriops clusia* was selected as the base genome to produce the hydrophiloid base genome optimized probe set, which after paralogy filtering targets 1,153 loci using 13,636 probes. This probe set will hereafter be referred to as the Hydro 1kv1.

**Fig. 2.**
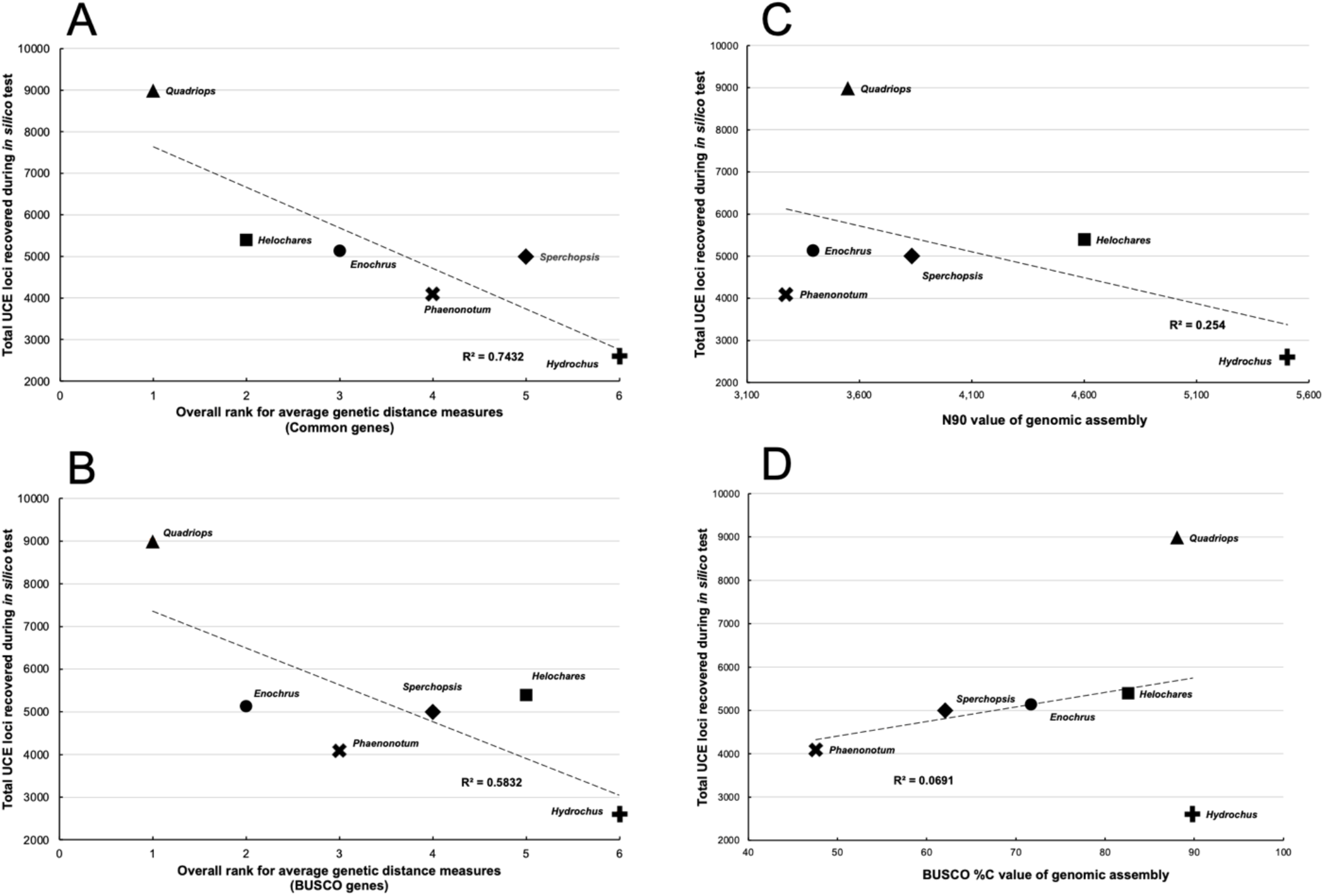
UCE loci recovery plotted against different metrics: A) average genetic distance estimated based on common gene fragments used in phylogenetics; B) average genetic distance estimated using BUSCO genes; C) N90 assembly metrics; and D) BUSCO %C values.

The Scarabaeidae base genome trials similarly revealed different base genomes resulted in considerable differences in the number of loci recovered (Table S19). The highest total loci recovered was 12% higher than the second highest total using a different genome. The *Onthophagus taurus* genome (Table 1) produced the third highest total recovered loci out of the five base genomes. For the reasons discussed above, the NCBI *Onthophagus taurus* genome was selected to serve as the base genome to produce the first Scarabaeid probe set, which after paralogy filtering targets 1,521 loci using 15,139 probes. This probe set will hereafter be referred to as the Scarab 1kv1.

### Optimization of initial design parameters

Changing the number of taxa the base genome must share a locus with for identification of putative loci from all to just one, dramatically increased the number of recovered loci during *in silico* testing. For the Hydrophiloid probe design, optimizing initial design parameters in this way significantly increased locus recovery (*P* = 4.794e^-5^ paired, two-tailed t-test on number of loci recovered in each taxon) and resulted in 85% more total loci recovered. Using both the optimal base genome of *Quadriops clusia* and initial design parameters of +1 for putative locus identification, resulted in a probe set which after paralogy filtering targets 1,886 loci using 22,341 probes. This probe set will hereafter be referred to as the Hydro 1.8kv1.

The Scarabaeidae probe design showed similar results where optimizing initial design parameters also significantly increased locus recovery (*P* = 2.764e^-6^ paired, two-tailed t-test on number of loci recovered in each taxon), but with a far more substantial increase in the total number of loci recovered at 205% (tripling the total). With *Onthophagus taurus* as the base genome and initial design parameters set to +1 for putative locus identification, after paralogy filtering this probe set targeted 2,343 loci using 23,278 probes. However, three loci could not be identified within the *Onthophagaus taurus* genome and were removed, leaving 2,340 loci and 23,260 probes. This probe set is dubbed the Scarab 2kv1.

### Combining probe sets

When the entire Coleoptera 1.1kv1 probe set was applied to our hydrophiloid taxa *in silico*, 957 loci could be recovered. We thus whittled the Coleoptera 1.1kv1 probes down to only these loci using the R script whittle_uce_probes.R (Alexander 2018), prior to merger with the fully optimized Hydro 1.8kv1 probe set. Of the 1,886 loci identified through the *de novo* Hydro 1.8kv1 probe design, only 82 were found in common with the 957 loci from the Coleoptera 1.1kv1 during merger. The final Hydrophiloid probe set, Hydro 2.7kv1, targets 2,761 loci (1,804 tailored, 957 generalized) using 32,657 probes.

*Onthophagus taurus* was included in the Coleoptera 1.1kv1 probe set design (Fig. 1), and as such has probes tailored to it. This design also utilized a strepsipteran, *Mengenilla moldrzyki*, as an outgroup for highly conserved locus identification (Fig. 1) (Faircloth 2017). Therefore, for the final Scarabaeidae probe set, using PHYLUCE we limited the Coleoptera 1.1kv1 probes to only those designed with *Onthophagus* and *Mengenilla* prior to merger. Of the 2,340 loci identified through the *de novo* Scarab 2kv1 probe design, 228 loci were in common with the 1,062 loci of the Coleoptera 1.1kv1 design. The final Scarabaeid probe set, Scarab 3kv1, targets 3,174 loci (2,112 tailored, 1,062 generalized) using 25,786 probes.

### Identification of loci

Use of *Onthophagus* as the base genome for the Scarab probe sets allowed all loci to be identified (Fig. 3, Table S21). We were also successful in identifying slightly over half of the loci associated with the final hydrophiloid probe set (Fig. 3, Table S21). Comparison of the proportions of different types of loci shows the majority are exonic, followed by intronic, then intergenic, with few being both intronic and exonic (Fig. 3). Of the identifiable loci in the Hydro 2.7kv1 probe set, the same trends are evident (Fig. 3). Optimization of initial design parameters increased the proportion of exonic loci, as did merging the probe set tailored probe set with the existing generalized probe set (Fig. 3).

**Fig. 3.**
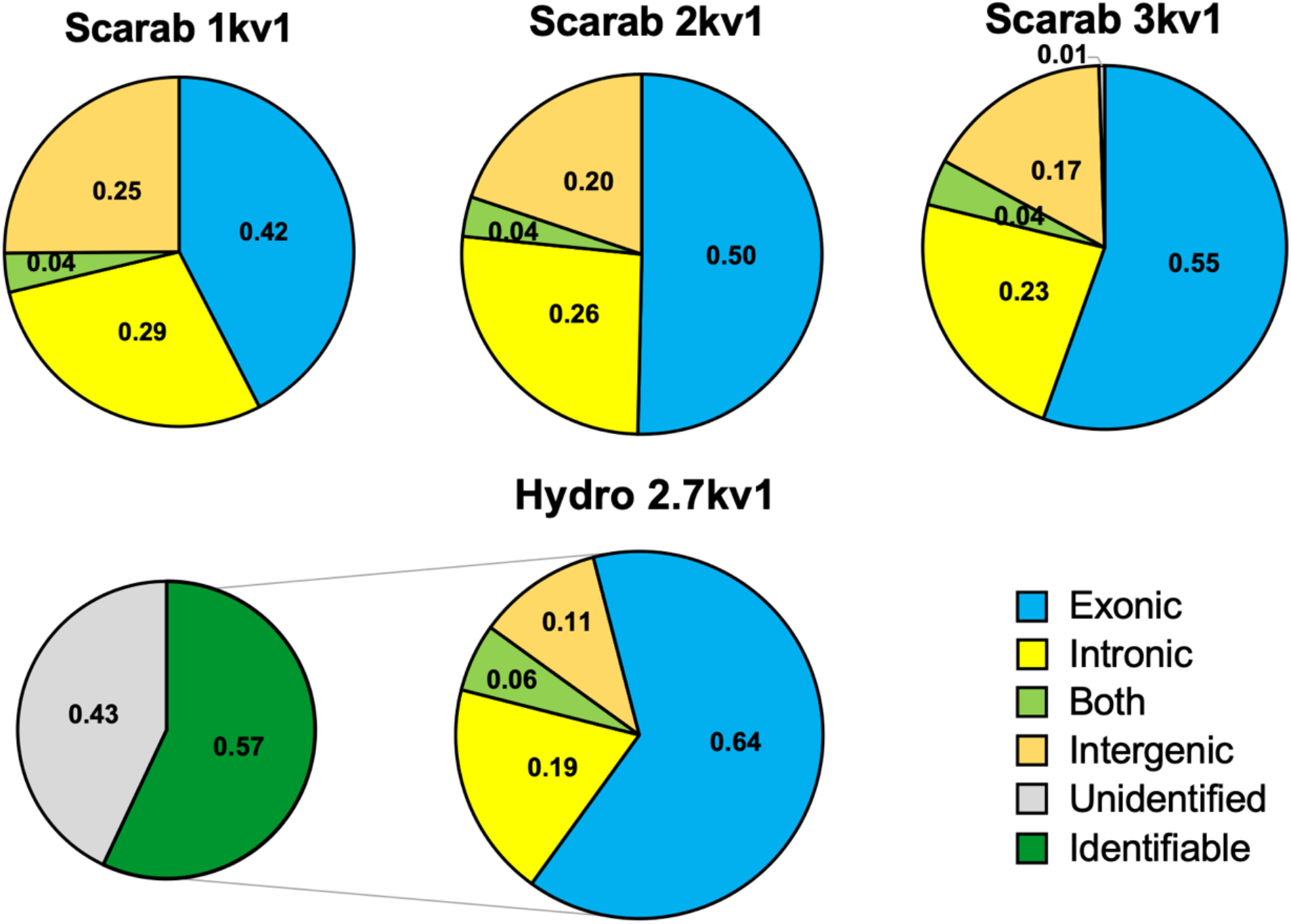
The proportion of different types of loci targeted by scarab and hydrophiloid probe sets.

### Comparison of phylogenetic utility of the probe sets

The probe set used had major impacts on aspects of the 75% complete data matrix (hereafter 75CM) assembled (Table 2). Use of any of the Scarab probe sets, which are phylogenetically closest to the scarabaeoid taxa (Fig. 1), resulted in the largest alignments with the highest proportions of both variable- and parsimony informative sites. Phylogenetic analysis of the 75CM produced with the Scarab 3kv1 probe set, resulted in a fully resolved tree whose topology was almost completely maximally supported (Figs. 4A,S1). The other scarab probe sets produced trees most similar to that shown in (Figs. 4A, S2–3).

**Table 2.** Results of the *in silico* test of different probe sets in terms of the 75% complete unpartitioned concatenated matrix and Robinson-Foulds distance of the resultant phylogenetic tree.

| Probe design | Alignment length<br>(bp) | Proport.<br>variable sites | Parsimony<br>informative sites<br>(bp) | Proport. parsimony<br>informative sites | Sum of plain<br>pair-wise R-F<br>distances |
| --- | --- | --- | --- | --- | --- |
| Adephaga 2.9kv1 | 54,189 | 0.421 | 19,549 | 0.361 | 52 |
| Coleoptera 1.1kv1 | 67,806 | 0.384 | 21,562 | 0.318 | 36 |
| Hydro 1kv1 | 25,798 | 0.417 | 8,533 | 0.331 | 36 |
| Hydro 1.8kv1 | 39,584 | 0.431 | 13,592 | 0.343 | 28 |
| Hydro 2.7kv1 | 157,675 | 0.422 | 55,745 | 0.354 | 26 |
| Scarab 1kv1 | 148,870 | 0.455 | 54,944 | 0.369 | 26 |
| Scarab 2kv1 | 214,531 | 0.465 | 81,820 | 0.381 | 24 |
| Scarab 3kv1 | 267,822 | 0.450 | 98,823 | 0.369 | 24 |

**Fig. 4.**
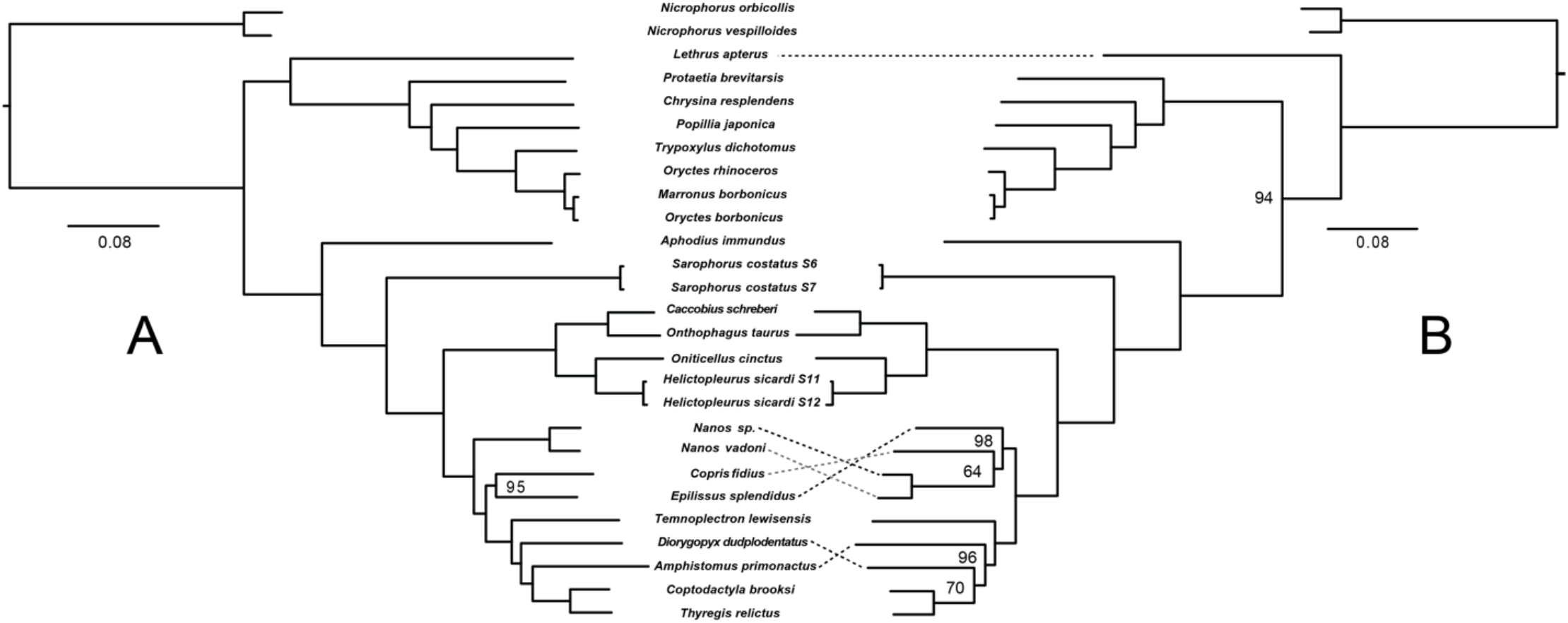
Phylogenetic trees of Scarabaeidae generated using UCEs; node values indicate bootstrap support. A) The tree produced with the Scarab 3kv1 probe set. B) The topology produced with the Adephaga 2.9kv1 probe set.

Among Hydrophiloidea probe sets (Table 2), representing the next closest clade phylogenetically to the Scarabaeoidea (Fig. 1), the Hydro 2.7kv1 produced a 75CM comparable to that of the Scarab 1kv1 and an identical phylogenetic tree (Figs. S3 & S4). On the contrary, the Hydro 1- and 1.8kv1 probe sets consisting solely of probes tailored for Hydrophiloidea, produced the shortest alignments with relatively low proportions of proportions of parsimony informative sites. The Hydro 1.8kv1 (Fig. S5) produced trees closer to the scarab probe sets than others, but Hydro 1kv1 produced a tree (Fig. S6) 10 R-F distances points away from both the Hydro 2.7 and Scarab 1kv1 (Table 2).

The Coleoptera 1.1kv1 was developed as a generalized or ‘universal’ probe set for Coleoptera, however, genomes used in locus identification for probe design consisted entirely of members of Polyphaga, the suborder of beetles Scarabaeoidea belongs to (Fig. 1). Therefore, this probe set represents the penultimate in terms of phylogenetic distance. The 75CM produced (Table 2) was larger in alignment length than both tailored hydrophilid probe sets, but had the overall lowest proportions of variable- and parsimony informative sites. The phylogenetic tree produced (Fig. S7) was equidistant with the Hydro 1kv1 from the Scarab 3kv1.

The probe set most phylogenetically distant from Scarabaeoidea is the Adephaga 2.9kv1, which was tailored for use in an entirely different suborder of beetles (Fig. 1). This probe set produced the most different tree (Figs. 4B, S7) as indicated by the largest R-F distance from all other trees (Table 2). The 75CM produced an alignment shorter than the generalized Coleoptera 1.1kv1, but longer than both tailored Hydro probe sets and with higher proportions of parsimony informative sites than all three (Table 2).

### Comparison of resulting phylogenetic trees

In general, the two contrasting phylogenetic trees for Scarabaeidae generated from different probe sets (Fig. 4A,B), are in agreement with previous studies. A clade in which Dynastinae is nested within Rutelinae emerges as sister to Cetoniinae, while Scarabaeinae is recovered as sister to Aphodiinae (Ahrens et al. 2019, McKenna et al. 2019). The scarabaeine genus *Sarophorus* is found to be the sister lineage to the rest of Scarabaeinae; Onticiellini+Onthophagini and Autralasian scarabaeines are recovered to be monophyletic; two Madagascan lineages, represented by the genera *Nanos* and *Epilissus*, come out as distantly related (Gunter et al. 2016, Tarasov and Dimitrov 2016).

However, some differences were observed between the trees generated using different probe sets. Specifically, within Scarabaeidae, the position of *Lethrus apterus* (Geotrupidae) alternated between being at the base of all Scarabaeidae (Fig. 4B), or at the base of the Pleurostict scarabs (Fig. 4A). Recent studies have supported the former relationship, which appears more plausible (Ahrens et al. 2019, McKenna et al. 2019). Within Scarabaeinae, the position of *Nanos* in relation to *Copris* and *Epilissus*, and *Amphistomus* and *Diorygopyx* within the Australasian Endemic clade varied. Considering that the present analysis was limited to taxa with genomes, it should be noted that differences were generally to be expected. Consequently, we anticipate that these discrepancies will be resolved once the taxon sampling is expanded.

## Discussion

### Identification and importance of selecting an optimal base genome

Our study confirms the previous findings of Gustafson et al. (2019) that base genome choice significantly alters probe set performance, based on *in silico* testing. Having an optimal base genome will result in the most loci recovered (Fig. 2A,B; Table S19), and based on comparisons of monolithic fasta files, loci that are longer than those developed using a less optimal base genome. Both factors are likely important for maximizing phylogenetic utility, as one will assemble matrices with larger alignment sizes through the increased locus recovery, and the longer size of these loci likely afford more variable sites for analysis.

In regards to how researchers interested in developing their own UCE probe sets can identify an optimal base genome efficiently, our results support Gustafson et al.’s (2019) finding that base genome performance negatively correlates with overall genetic distance away from taxa used in probe design (Fig. 2A,B). If highly complete genome assemblies are available, genes extracted as part of BUSCO assessment can be used to estimate genetic distances, or gathering traditional Sanger loci used in phylogenetic studies provides similar results (Fig. 2A,B). Our results also corroborate that genome assembly metrics do not correlate with base genome performance (Fig. 2C,D). Thus, genome assembly completeness alone, is not a good justification for selecting a taxon to serve as the base genome, as this does not guarantee optimal probe design.

### Relaxed initial locus identification provides more exonic loci

Decreasing the number of taxa candidate loci must be shared between to only one during initial probe design dramatically increased the total number of loci UCE loci recovered (Table S20), another result consistent with Gustafson et al. (2019). Furthermore, after filtering paralogous loci, the additional loci targeted appear to be skewed towards exonic regions, based on the evident increase in the ratio of exonic loci between the Scarab 1kv1 and Scarab 2kv1 (Fig. 3). Investigators developing their own UCE probe sets wanting to increase the number of loci targeted, and those interested in maximizing exonic loci for analysis, should consider relaxed initial locus identification with candidate loci required to be shared between the base genome and just one other taxon. Under this scenario, stringent paralogy filtering is recommended, which can scarabaeoid be accomplished using the scripts available from Alexander (2018).

### Combining generalized- and tailored probe sets

Combining the generalized probes from the Coleoptera 1.1kv1 (Fig. 1) with tailored hydrophilid and scarab probes improved phylogenetic utility of final probe sets via increased alignment length and higher proportion of parsimony informative sites in the 75CM (Table 2). The addition of these probes also had the effect of decreasing the proportion of variable sites. The Coleoptera 1kv1 probe set was developed to target loci across highly divergent lineages of beetles (Fig. 1), and as a result, the loci its probes pull contain decreased variability. While this aspect may seem like a potential drawback, a mixture of highly conserved loci and more variable tailored loci has been shown to improve phylogenetic analysis, especially in regards to placement of phylogenetically distant taxa (Gustafson et al. 2020). Consistent with this, the Hydro 2.7kv1, which contains generalized probes, outperformed both tailored-only hydrophiloid probe sets when applied to the distantly related scarabaeoid taxa (Table 2).

An additional important aspect of combining probe sets is affording cross compatibility between probe sets and phylogenetic studies utilizing them (Fig. 1). Thus, UCE enriched read data generated using the two novel probe sets developed here, will share targeted loci and be combinable with those from studies utilizing the Coleoptera 1.1kv1 like Kobayashi et al. 2021, Bradford et al. 2022, Sota et al. 2022, etc.; and to a lesser extent those using the Adephaga 2.9kv1 probe set (Fig. 1).

### Selecting an existing probe set for UCE phylogenomics

Predictably, as evolutionary distance of taxa used in probe design increased away from Scarabaeoidea (Fig. 1), phylogenetic utility of probe sets decreased, as evidenced in smaller alignment lengths, decreased proportions of variable- and parsimony informative sites, and lower numbers of parsimony informative sites of the 75CM (Table 2). Furthermore, trees produced using probe sets further away from Scarabaeoidea received lower support (Figs. S1–S8) and had more strongly differing topologies (Table 2, Figs. S1–S8). Notably, taxonomic depth of probe design also had a strong impact. The overall worst performing probe set (Table 2) was not the Adephaga 2.9kv1, the furthest away phylogenetically (Fig. 1), but the Hydro 1kv1 probe set tailored for use in the hydrophiloid clade (Fig. 1). The reason probe sets containing only tailored hydrophiloid probes performed in the most limited fashion, likely has to do with locus identification being limited only to the focal clade Hydrophiloidea (Fig. 1). Thus, many of these loci are unlikely to be present, or at least conserved enough, at deeper levels in the Coleoptera tree, including within Scarabaeoidea, for UCE probes to recover them. Optimizing initial locus identification improved phylogenetic performance in the tailored-only Hydro 1.8kv1 (Table 2), especially in regards to the tree produced, but the 75CM was still limited relative to the general Coleoptera 1.1kv1. This latter probe set was meant to serve as a generalized Coleoptera probe set (Fig.1), and as such it provides conserved loci which the lowest proportions of variable sites, but ultimately functions better than probe sets tailored for specific clades outside of our focal Scarabaeoidea (Table 2). Notably, combining the generalized probes Coleoptera probes with the optimized hydrophiloid tailored probes in the Hydro 2.7kv1 expanded the probe set’s phylogenetic utility to comparable to a incompletely optimized scarab probe set like the Scarab 1kv1 (Table 2).

Given the above, when selecting an existing probe set for a focal clade, the primary concern is phylogenetic distance (Fig. 1). All scarab probe sets performed the best on scarabaeoid taxa during *in silico* testing (Table 2). However, care should be taken to ensure the probe set includes some probes developed with taxonomic breadth beyond a highly derived focal clade, as tailored-only probes for the sister clade outside of scarabaeoids (i.e., the Hydro 1kv1 and 1.8kv1, Fig. 1) performed comparable to- or worse than a generalized only probe set like the Coleoptera 1.1kv1 (Table 2). For example, using Fig. 1, researchers interested in selecting an existing probe set for use within Staphylinoidea, the Scarab 3kv1 probe set is likely to perform the best given Scarabaeidae is sister to Staphylinoidea and includes generalized probes. Because the Hydro 2.7kv1 also includes generalized probes, it would likely function better than picking the existing Coleoptera 1.1kv1 alone, but our results suggest not as well as the Scarab 3kv1. For researchers interested in selecting an existing UCE probe set for broader use within Curculionidae (Fig. 1), unless the study is focused on Entiminae, the choice is less clear. While the Pachyrhynchini probe set of Van Dam et al. (2022b) was developed specifically for a tribe of entimine weevils and thus is the phylogenetically closest probe set available, suggesting potential optimal performance, there is the possibility that its design for a highly derived clade could limit its utility in the absence of generalized probes. A possible solution would be to include all *Dendroctonus* probes from the Coleoptera 1.1kv1 and the entire Pachyrhynchini set, but the size of this probe set: 225,632 probes targeting 12,522 (Van Dam et al. 2022b) may prohibit its combined use, as third party sequencing companies often have price increases associated with higher number of probes synthesized.

## Conclusion

Our study confirms previous recommendations for developing an optimized UCE probe design. Selecting a taxon to serve as the base genome that has the shortest overall genetic distance from all other taxa involved in probe design, and setting the initial locus identification parameters to loci shared between this base genome and just one other taxon, will result in an optimized probe set that recovers the most loci during *in silico* testing, and which have the longest per-locus alignments. We also found that genome assembly metrics, such as completeness measures, do not correlate well with base genome performance, and alone are not a good justification for selecting a taxon to serve as the base genome. However, use of an annotated genome as the base will allow identification of UCE loci targeted by the probe design, which can in turn be used to merge cogenic loci for analysis, and prevent development of off-target probes, both of which are important aspects to consider (Van Dam et al. 2021, 2022a). Finally, we found further evidence that combining generalized probes with tailored probes improves phylogenetic performance of the probe set. Applying these aspects to probe design we developed two novel beetle probe sets: the Hydro 2.7kv1 targeting 2,761 loci (1,804 tailored, 957 generalized) using 32,657 probes; and the Scarab 3kv1 targeting 3,174 loci (2,112 tailored, 1,062 generalized) using 25,786 probes. Both are available under the public domain license (CC-0) and can be accessed.

While developing tailored UCE probes for focal taxa appears to be the best method to ensure optimal *in vitro* UCE locus recovery, selecting an existing probe set for use may be necessary. Our results suggest that when selecting among UCE probe sets, those phylogenetically closest to the ingroup are likely to perform the best. Additionally, probe sets containing probes developed across a broader taxonomic breadth, in addition to tailored probes for the focal clade, are likely to perform better than selecting a ‘universal’ probe set consisting solely of generalized probes. We also found that probe sets consisting solely of probes tailored to a more derived clade, phylogenetically adjacent to focal taxa, may perform worse than generalized probes alone. Ultimately, all *in silico* testing, including that conducted here, should be considered exploratory, requiring *in vitro* validation, as *in vitro* results differ for a variety of reasons including physical probe synthesis, probe efficacy, and stochastic error. While our study has focused on beetles, we expect our findings to be broadly applicable to arthropod UCE probe design and application, given UCEs in both groups are primarily exonic (Van Dam et al. 2021) and its members possess broad genomic- and taxonomic diversity.

## Supplementary Materials

The Supplementary Materials, including the designed UCE probe sets, is available at Zenodo (https://zenodo.org/record/7509385#.Y7f1d-xBwnc, DOI: 10.5281/zenodo.7509385.

## Author contributions

G.T.G, S.T. and N.L.G jointly conceived the project. G.T.G and R.D.G conducted analyses to design probes and performed simulation experiments. S.T. wrote the new R script. G.T.G and A.E.Z.S. devised, directed and funded the Hydrophiloidea component of the research. S.T and N.L.G devised, directed and funded the Scarabaeoidea component of the research. All authors discussed the results and contributed to the final manuscript.

## Acknowledgements

We are thankful to Giulio Montanaro and Michele Rossini for species identifications, and to Vasily Grebennikov and Geoff Monteith for providing specimens used in this study.

Computational analyses and data processing reported in this study were conducted on Northern Arizona University’s Monsoon computing cluster, funded by Arizona’s Technology and Research Initiative Fund and the University of Kansas Center for Research computer cluster. This material is based upon work supported by the US National Science Foundation under grants DEB-1942193 to NLG and DEB-1453452 to AEZS. The work of ST was supported by the Academy of Finland grant: 331631, and three-year grant from the University of Helsinki.

